# Inducible TRAP RNA profiling reveals host genes expressed in Arabidopsis cells haustoriated by downy mildew

**DOI:** 10.1101/2023.01.17.524393

**Authors:** Shuta Asai, Volkan Cevik, Jonathan D.G. Jones, Ken Shirasu

## Abstract

The downy mildew oomycete *Hyaloperonospora arabidopsidis*, an obligate filamentous pathogen, infects Arabidopsis by forming feeding structures called haustoria inside host cells. Previous transcriptome analyses revealed host genes are specifically induced during infection; however, RNA profiling from infected tissues may fail to capture key transcriptional events occurring exclusively in haustoriated host cells where the pathogen injects virulence effectors to modulate host immunity. To determine interactions between Arabidopsis and *H. arabidopsidis* at the cellular level, we devised a new translating ribosome affinity purification system applicable to inducible, including pathogen-responsive, promoters thus enabling haustoriated cell-specific RNA profiling. Among the host genes specifically expressed in *H. arabidopsidis*-haustoriated cells, we found genes that promote either susceptibility or resistance to the pathogen, providing new insights into the Arabidopsis/downy mildew interaction. We propose that our novel protocol for profiling cell-specific transcripts will be applicable to several stimulus-specific contexts and other plant-pathogen interactions.

## Introduction

*Hyaloperonospora arabidopsidis* causes downy mildew disease in the model plant Arabidopsis. *H. arabidopsidis* is an obligate biotrophic oomycete that completes its life cycle without killing the host. Asexual *H. arabidopsidis* conidiospores geminate and form appressoria to penetrate leaf surfaces. Hyphae then grow intercellularly, producing numerous pyriform-shaped structures called haustoria in mesophyll cells (Coates and Beynon, 2010). Haustoria impose invaginations of the plant cell, creating an interface between host and pathogen called an extra-haustorial matrix. This matrix is thought to be the site where the pathogen acquires nutrients from the plant and where pathogen-derived effectors are delivered into the host cell to suppress defense responses and promote susceptibility.

Host genes that promote susceptibility to pathogens are called susceptibility (*S*) genes (van Schie and Takken, 2014). *S* genes are generally expressed in infected cells to accommodate pathogens. In the Arabidopsis/downy mildew interaction, for example, the *S* gene *DMR6 (Downy Mildew Resistant 6*) is exclusively induced in host cells containing haustoria (haustoriated cells) (Fig. 1A, van Damme et al., 2008). *DMR6* encodes a salicylic acid (SA) 5-hydroxylase that inactivates SA, a phytohormone essential for plant immunity (Zhang et al., 2017). Consistently, *H. arabidopsidis* specifically suppresses SA-inducible *PR1 (PATHOGENESIS-RELATED GENE1*) expression in haustoriated cells, whereas *PR1* is expressed in the surrounding cells (non-haustoriated cells) (Fig. 1A, Caillaud et al., 2013; Asai et al., 2014). Several *H. arabidopsidis* effectors are able to suppress the SA signaling pathway (Caillaud et al., 2013; Asai et al., 2014; Wirthmueller et al., 2018); however, little is known about what events occur in the infected cells to modulate the local responses of Arabidopsis to *H. arabidopsidis*. Identifying these events requires cell-specific transcript analysis.

**Figure 1.**
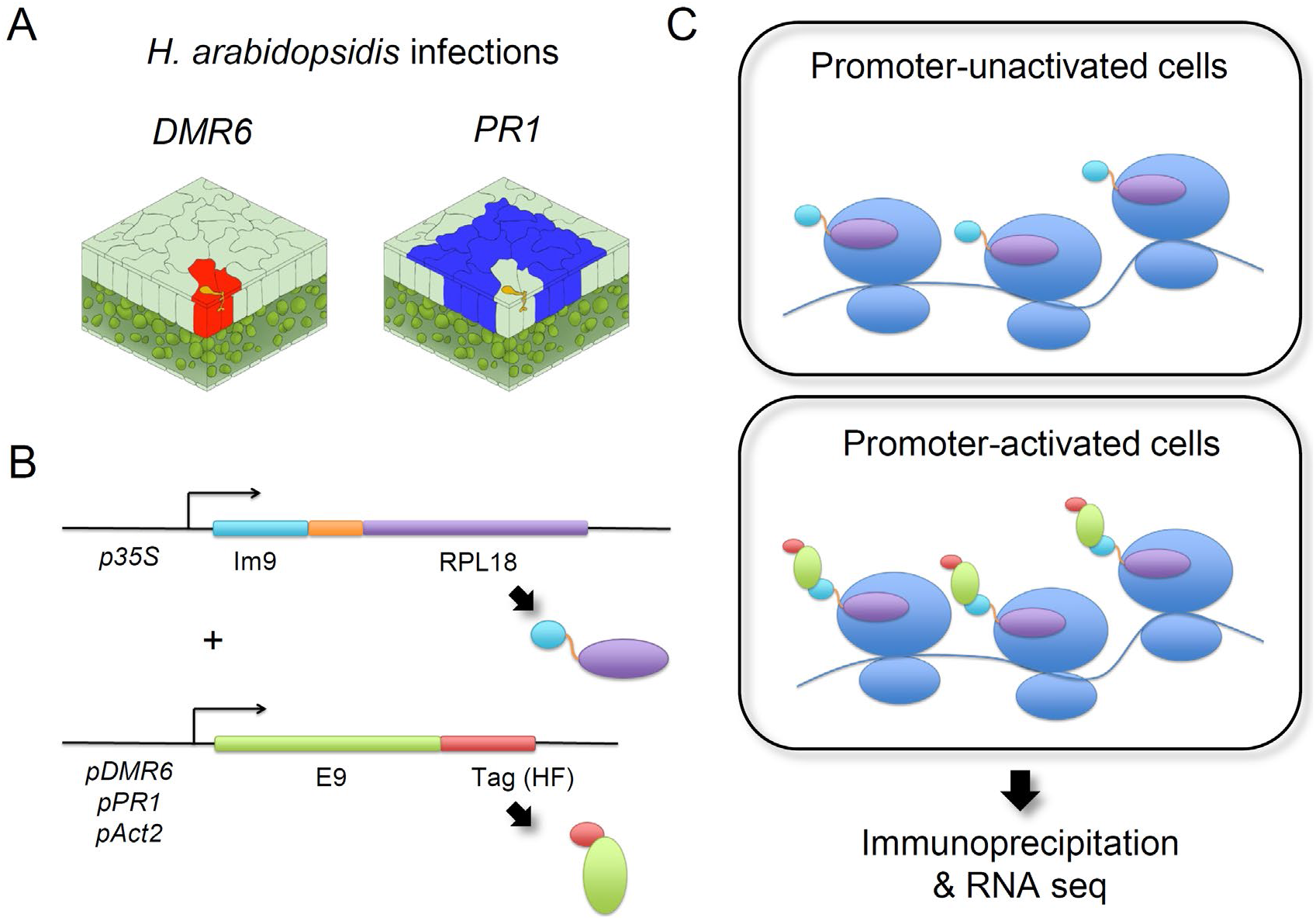
Schematic diagram of a new translating ribosome affinity purification (TRAP) system. (**A**) Schematic view of cell-specific responses in the *H. arabidopsidis–* Arabidopsis interaction. *H. arabidopsidis* extends hyphae to form haustoria inside host cells (yellow shapes). Red-shaded cells indicate cells in which the *DMR6* promoter (*pDMR6*) is activated, i.e., the haustoriated (infected) cells. Blue-shaded cells indicate cells in which the *PR1* promoter (*pPR1*) is activated, i.e., the non-haustoriated adjacent (non-infected) cells. (**B**) Schematic representation of two chimeric constructs; Im9-RPL18 fused to the *35S* promoter (*p35S*) and E9-HF controlled by *pDMR6, pPR1*, or the *Actin2* promoter (*pAct2*). HF, a tandem 6xHis and 3xFLAG epitope tag. (**C**) Schematic diagram of ribosomal complexes in cells where the promoters fused to E9-HF are unactivated (upper panel) or activated (lower panel).

Translating ribosome affinity purification (TRAP) is a powerful method that enables cell type-specific RNA profiling (Mustroph et al., 2009b; Heiman et al., 2014). In the traditional TRAP system, ribosome-associated mRNAs are immunopurified from specific cell populations that express an epitope-tagged ribosomal protein via developmentally regulated promoters (i.e., cell type-specific promoters) (Fig. S1) (Mustroph et al., 2009b). A limitation of the traditional TRAP methodology makes the procedure inapplicable to cells in which stress-responsive promoters are activated because the newly synthesized epitope-tagged ribosomes must replace pre-existing ribosomes in the cells, i.e., a problem of ribosomal turnover. To overcome this limitation, an affinity tag, but not a ribosomal protein, should be controlled by the specific promoter to capture ribosomes with corresponding tags under the control of their own or a constitutive promoters. Based on this concept, we established a novel TRAP system that relies on high affinity colicin E9-Im9-based interactions. Our new system allows the formation of tagged ribosomal complexes only in cells where the *DMR6* promoter is activated, thereby enabling haustoriated cell-specific RNA profiling. Among the haustoriated cell-specific transcripts, we found genes involved in resistance and susceptibility to *H. arabidopsidis*, indicating that haustoriated cell-specific RNA profiling can provide new insights into the interaction between Arabidopsis and the downy mildew pathogen.

## Results

### A new TRAP system for cells with specific promoter activation

Although *DMR6* and *PR1* show distinct cellular expression patterns in Arabidopsis infected with *H. arabidopsidis* (Fig. 1A; van Damme et al., 2008; Caillaud et al., 2013; Asai et al., 2014), transcriptome analysis using whole tissues revealed no significant difference in the expression patterns of these genes during infection (Fig. S2; Asai et al., 2014). To elucidate the interaction between Arabidopsis and *H. arabidopsidis* at the cellular level, we designed a new TRAP system using two high-affinity binding proteins: a bacterial toxin protein, E9, and its cognate immunity protein, Im9 (Li et al., 1997). The new TRAP system consists of two chimeric transgenes: one gene encodes RPL18 fused to Im9 driven by the *35S* promoter (*p35S);* the second gene is controlled by promoters of stress-responsive genes such as *DMR6* (*pDMR6*) or *PR1* (*pPR1*) and encodes E9 fused to a tandem 6x<u>H</u>is and 3x<u>F</u>LAG epitope tag (HF) used for purification (Fig. 1B). In cells where the corresponding promoters are active, the purification tag attaches to ribosomes when binding between E9 and Im9 occurs (Fig. 1C).

We confirmed whether tagged ribosomes are formed by the binding of E9 and Im9 using a *Nicotiana benthamiana* transient expression system. As expected, YFP-RPL18 accumulated in the nucleolus, where most ribosome biogenesis events take place (Fig. 2). E9-GFP localized to the cytoplasm and nucleus, excluding the nucleolus, when coexpressed with GUS as a control, whereas GFP fluorescence was observed in the nucleolus when E9-GFP was coexpressed with Im9-RPL18 (Fig. 2). These results indicated that ribosomal complexes consisting of chimeric constructs were formed upon the binding of E9 and Im9.

**Figure 2.**
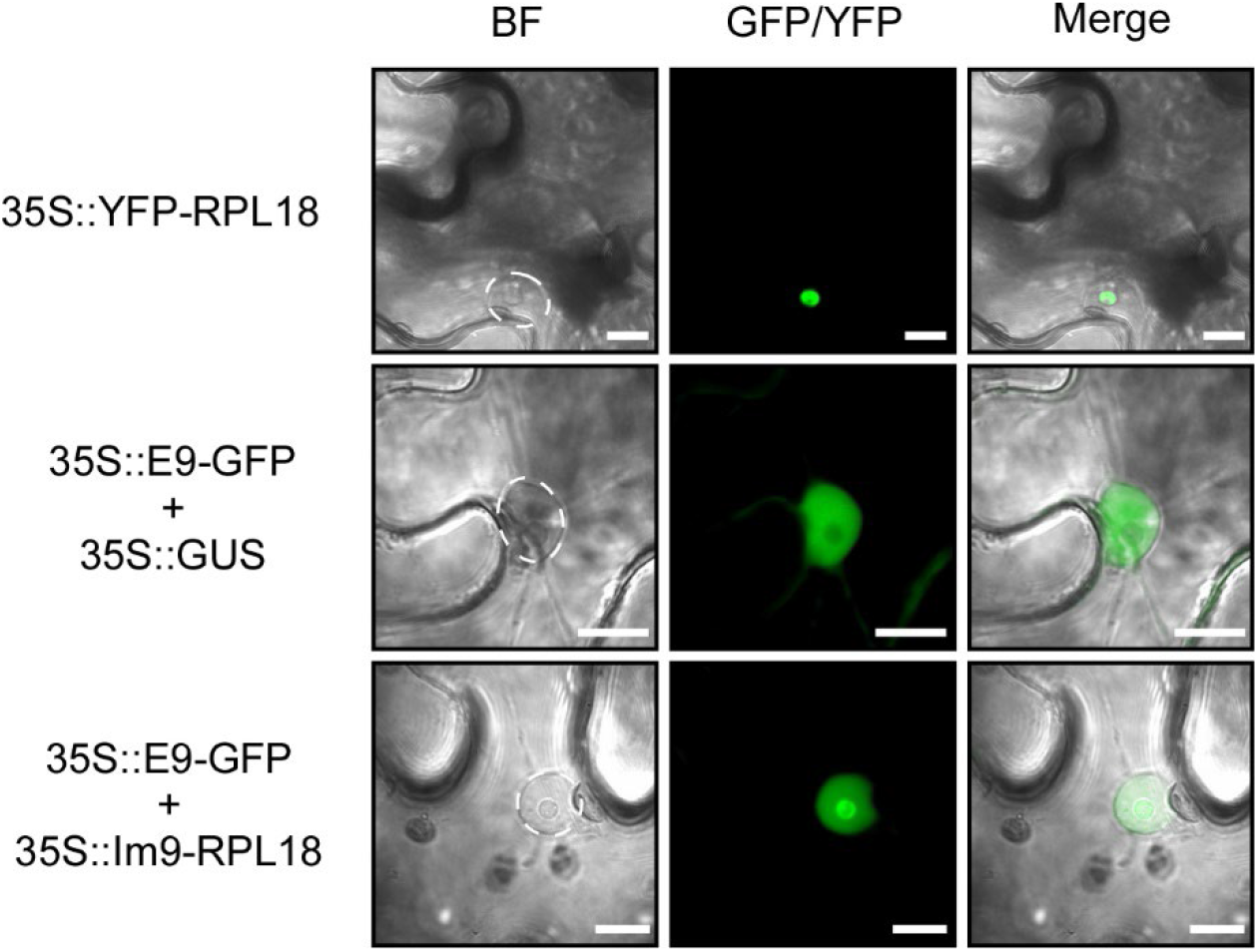
Formation of ribosomal complexes consisting of chimeric constructs coincident with E9 and Im9 binding. Subcellular localization of YFP-RPL18 and E9-GFP when coexpressed with GUS and Im9-RPL18. The indicated constructs were transiently expressed in *N. benthamiana* leaves. The left image is the bright-field (BF) image, the middle image is from the GFP/YFP channel, and the right image is the overlay of the BF image and GFP channel. Dashed white circles mark the locations of nuclei in the BF pictures. Scale bars, 10 μm.

### Validating the cell-specific TRAP system with *H. arabidopsidis*-infected Arabidopsis

We created Arabidopsis transformants containing two transgenes: Im9-RPL18 controlled by *p35S* (*p35S::Im9-RPL18*) and E9-HF driven by either *pDMR6* (*pDMR6::E9-HF*), *pPR1* (*pPRP::E9-HF*), or the *Actin2* promoter (*pAct2::E9-HF*) as a control (Fig. 1B). We hypothesized that E9-RPL18 would bind to Im9-HF in cells where both transgenes were expressed, thereby enabling conditional but efficient tagging of pre-existing ribosomes in the cells of interest (Fig. 1C). After inoculating the transformants with *H. arabidopsidis* virulent isolate Waco9, proteins derived from fractions containing ribosomes and mRNAs (polysome-enriched fractions, see Materials and methods) were extracted from infected tissues. The Im9-RPL18/E9-HF complexes were immunoprecipitated with anti-FLAG agarose beads, from which RNAs were extracted and referred to as RNAs_IP (Fig. 3A). We also extracted RNAs directly from the polysome-enriched fractions and designated those as RNAs_Total. To confirm whether E9-HF is properly controlled by *pDMR6* or *pPR1* in the transformants, immunoblots of protein samples after inoculation with *H. arabidopsidis* were probed with anti-FLAG antibodies. As expected, E9-HF was detected during *H. arabidopsidis* infection in transformants containing *pDMR6::E9-HF* or *pPR1::E9-HF*, whereas transformants containing *pAct2::E9-HF* constantly accumulated E9-HF (Fig. 3B). Importantly, RT-qPCR analysis confirmed that *DMR6* or *PR1* transcripts were enriched in the RNAs_IP samples derived from transformants containing *pDMR6::E9-HF* or *pPR1::E9-HF*, respectively, whereas the transcript levels of *Act2* were comparable among the RNAs_IP samples (Fig. 3C). In the RNAs_Total samples, there was no difference in the transcript levels of either *DMR6, PR1* or *Act2* (Fig. 3C). These results indicated that the new TRAP system successfully enriched specific cell-derived mRNAs during *H. arabidopsidis* infection.

**Figure 3.**
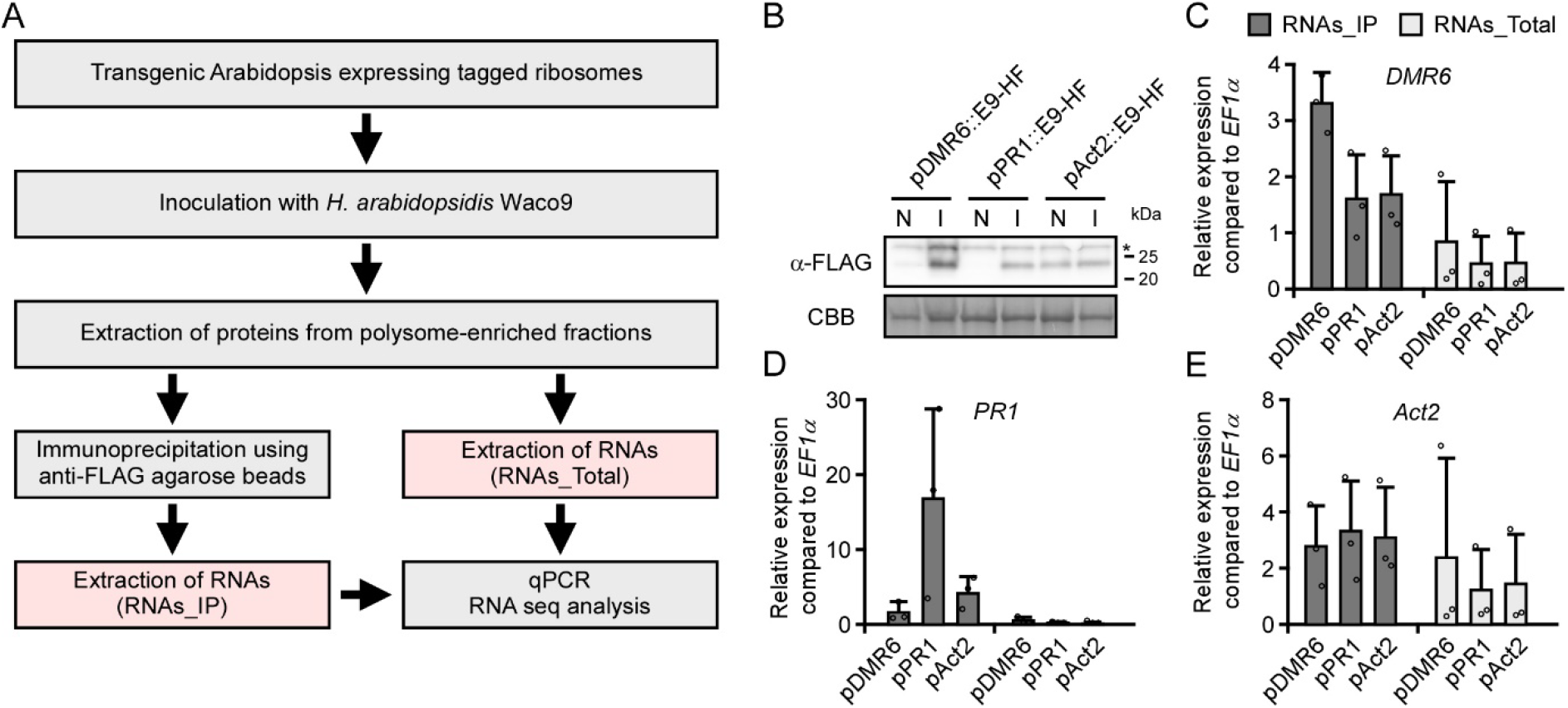
Validating the enrichment of specific cell-derived mRNAs during *H. arabidopsidis* infection by the new TRAP system. (**A**) Flowchart of the steps used to validate the cell-specific TRAP system. Protein accumulation (**B**) and expression of *DMR6* (**C**), *PR1* (**D**), and *Act2* (**E**) in Arabidopsis Col-0 transgenic lines containing *pDMR6::E9-HF* (pDMR6), *pPR1::E9-HF* (pPR1) or *pAct2::E9-HF* (pAct2) and *p35S::Im9-RLP18*. (**B**) Total proteins were prepared from 3-week-old plants at 5 d after spraying water (N) or inoculation with *H. arabidopsidis* (I). An immunoblot analyzed using anti-FLAG (upper panel) antibodies. Protein loads were monitored by Coomassie Brilliant Blue (CBB) staining of bands corresponding to ribulose-1,5-bisphosphate carboxylase (Rubisco) large subunit (lower panel). (**C-E**) The expression levels of *DMR6, PR1*, and *Act2* in the RNAs_Total and RNAs_IP samples were determined by RT-qPCR. Data are means ± SDs from three biological replicates.

### Identifying *DMR6-coexpressed* genes during *H. arabidopsidis* infection

To investigate cell-specific responses during *H. arabidopsidis* infection, the TRAP samples were subjected to RNA-seq analysis (Table S1). In the RNAs_Total samples, there were no differentially expressed genes in the *pDMR6::E9-HF* or the *pPR1::E9-HF* transformants compared to the *pAct2::E9-HF* control (Fig. 4A). By contrast, the RNAs_IP samples had genes with significant differences in expression levels (FDR = 0.05). The *pDMR6::E9-HF* transformants had 4,524 upregulated genes and 319 downregulated genes; whereas the *pPR1::E9-HF* transformants had 3,969 upregulated genes and 338 downregulated genes compared to the control (Fig. 4a and Table S2). Importantly, *DMR6* and *PR1* were among the upregulated genes of the *pDMR6::E9-HF* and *pPR1::E9-HF* transformants, respectively. To identify genes coexpressed with *DMR6* and are specifically expressed in cells infected by *H. arabidopsidis* (haustoriated cells), we compared the upregulated genes in the *pDMR6::E9-HF* transformants to those in the *pPR1::E9-HF* transformants. The comparison revealed 1,571 candidate genes coexpressed with *DMR6* but not *PR1* (Fig. 4B and Table S3). Candidate genes were further limited by a comparison with our previously reported list of genes whose expression was significantly upregulated during infection with *H. arabidopsidis* (Table S3; Asai et al., 2014). In this analysis, we identified *DMR6* and 53 genes that were designated *DMR6*-coexpressed genes (Table 1). Among these 54 genes, gene ontology (GO) analysis revealed an overrepresentation of genes related to disease resistance (e.g., GO:0050832 and GO:0006952) and genes responsive to oxygen levels (e.g., GO:0001666 and GO:0070482) and chemicals (e.g., GO:0042221) (Fig. S3).

**Figure 4.**
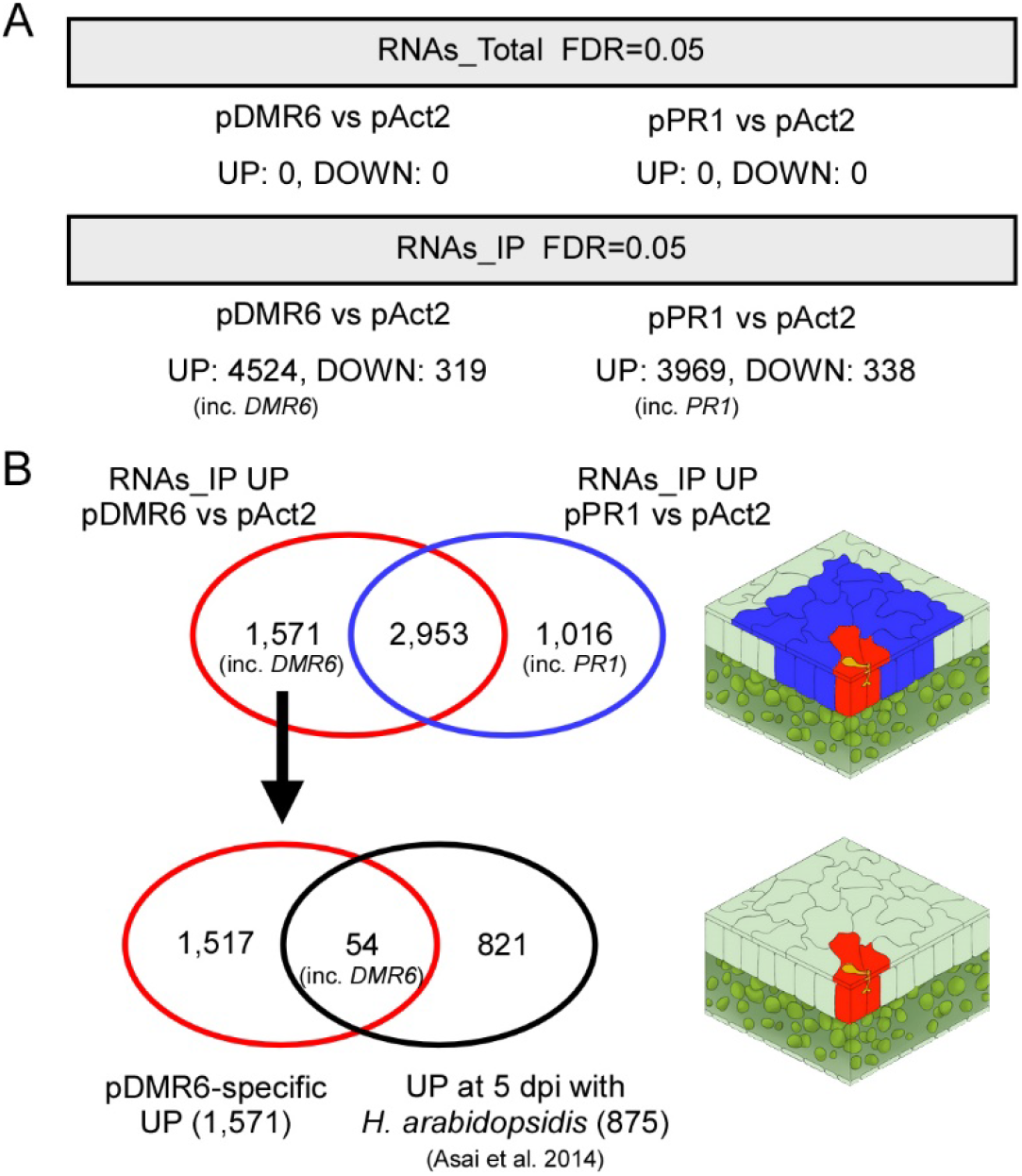
Selecting confident candidate *DMR6*-coexpressed genes. (**A**) The number of genes significantly upregulated (UP) or downregulated (DOWN) among Arabidopsis Col-0 transgenic lines containing *pDMR6::E9-HF* (pDMR6), *pPR1::E9-HF* (pPR1), or *pAct2::E9-HF* (pAct2) and *p35S::Im9-RLP18*. (**B**) Assessment of overlapping differentially expressed genes to select confident candidate *DMR6*-coexpressed genes. The comparison of upregulated genes between pDMR6 and pPR1 transformants in the RNAs_IP samples revealed 1,571 genes as *DMR6*-coexpressed candidate genes (pDMR6-specific UP). Comparing the 1,571 genes with 875 genes significantly upregulated at 5 d after inoculation (dpi) with *H. arabidopsidis* reported by Asai et al. (2014) revealed that 54 genes overlapped in the two conditions. The figures on the right indicate proposed expression sites: red-shaded cells, expression sites where *DMR6*-coexpressed genes are expressed; blue-shaded cells, expression sites where *PR1*-coexpressed genes are expressed.

**Table 1.**
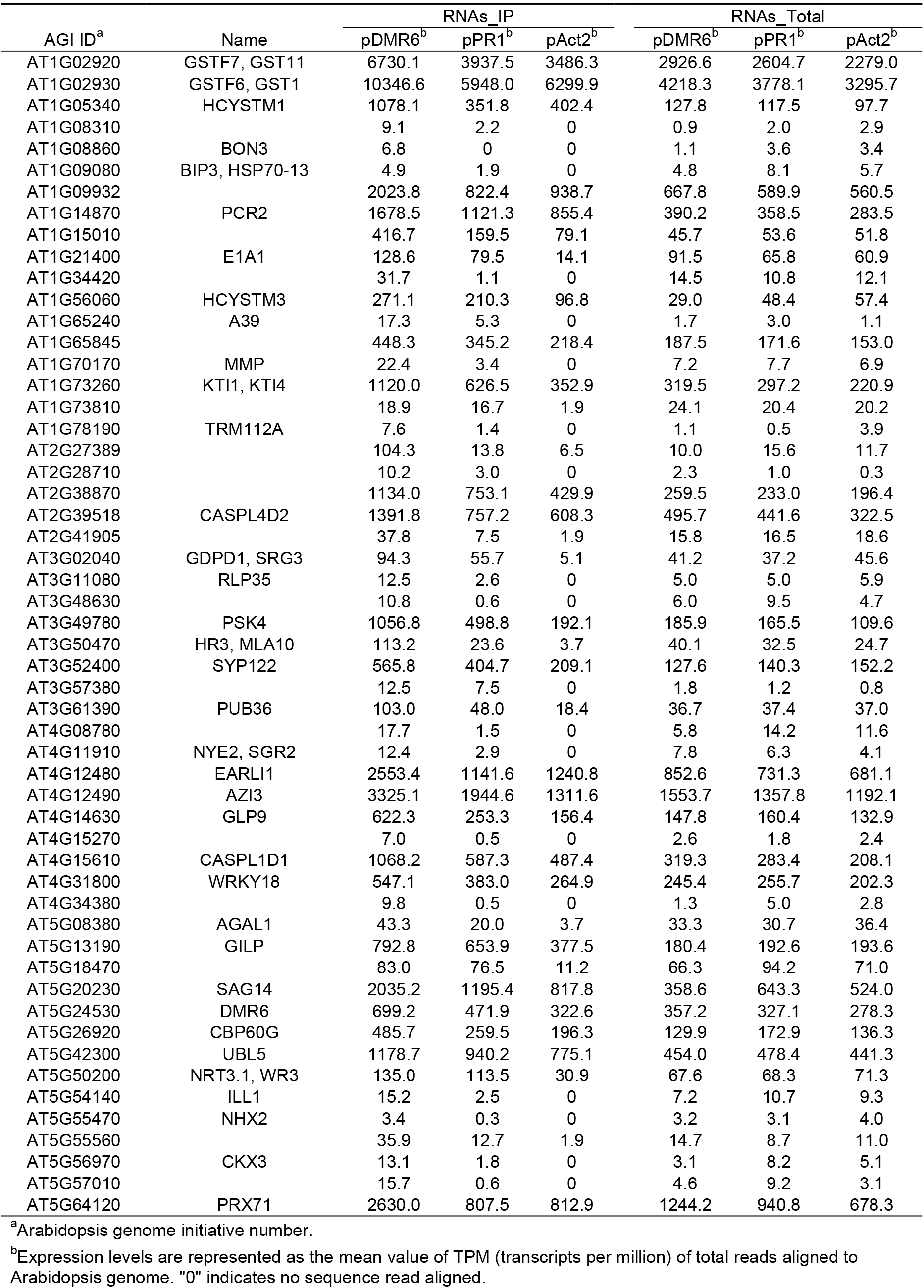
The list and expression patterns of *DMR6* and 53 *DMR6*-coexpressed genes.

In the *DMR6*-coexpressed gene list (Table 1), we found *PHYTOSULFOKINE 4 PRECURSOR* (*PSK4; AT3G49780*) and *WRKY18 (AT4G31800*), genes known to function as negative regulators of plant immunity. Arabidopsis transformants containing the *PSK4* or *WRKY18* promoter controlling the *GUS* reporter gene were generated and inoculated with *H. arabidopsidis* to confirm that *PSK4* and *WRKY18* are expressed in haustoriated cells. In both transformants, GUS staining was restricted to haustoriated cells as observed for *H. arabidopsidis*-infected *pDMR6::GUS* lines (Fig. 5). This result indicated that *PSK4* and *WRKY18* are expressed specifically in the cells haustoriated with *H. arabidopsidis*. These data also suggest that genes involved in plant immunity can be identified using the new TRAP system. Next, we randomly chose the following five genes from among the *DMR6*-coexpressed candidate genes (Table 1) for promoter-fused GUS analysis: *AZELAIC ACID INDUCED 3* (*AZI3; AT4G12490), KUNITZ TRYPSIN INHIBITOR 4* (*KTI4; AT1G73260*), *AT1G09932* (annotated to encode a phosphoglycerate mutase family protein), *PLANT CADMIUM RESISTANCE 2* (*PCR2; AT1G14870*), and *GERMIN-LIKE PROTEIN 9* (*GLP9; AT4G14630*). As expected, GUS staining was observed specifically in *H. arabidopsidis*-haustoriated cells in all transformants tested and in the *pDMR6*::*GUS* control (Fig. 5), indicating that these five genes are also coexpressed with *DMR6*.

**Figure 5.**
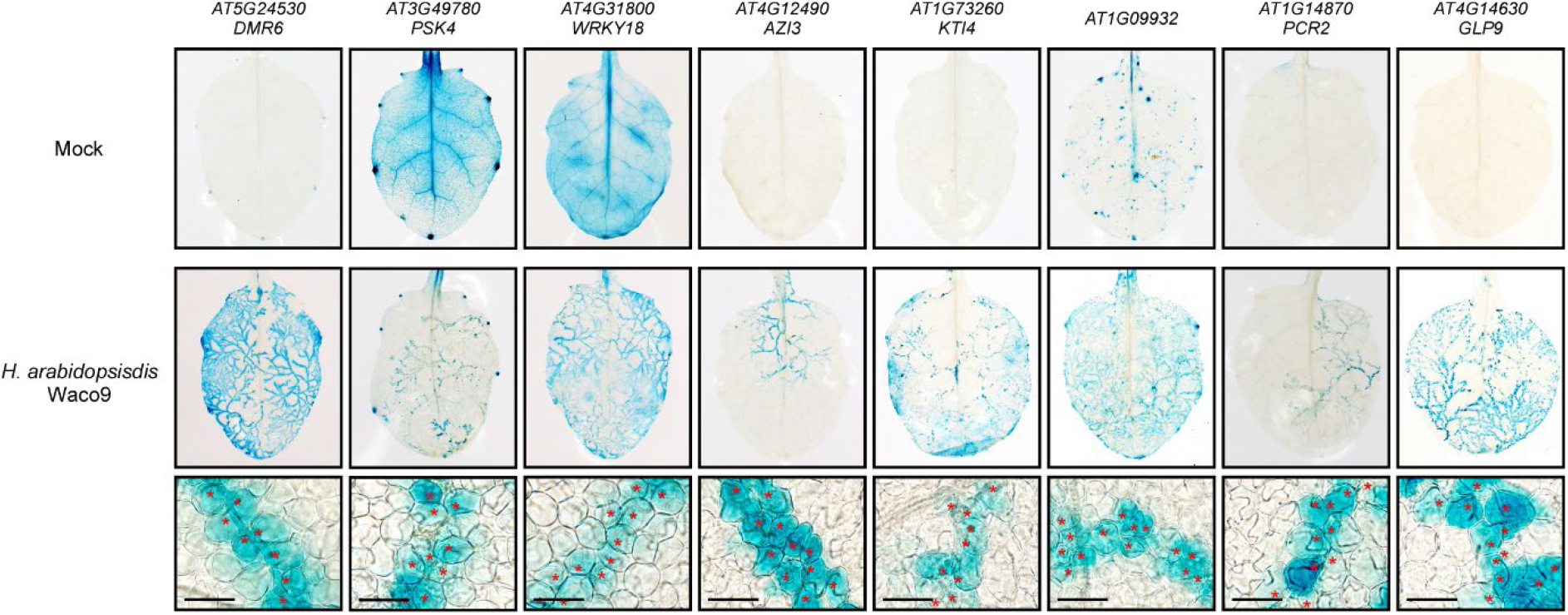
Cellular expression patterns of *DMR6*-coexpressed genes. GUS staining of 3-week-old Arabidopsis leaves containing the indicated gene promoter fused to a *GUS* reporter gene after inoculating leaves with *H. arabidopsidis* Waco9 and water as a control (Mock). A GUS staining solution containing one-fifth the amount of substrate was used to monitor expression in the infected leaves due to high promoter activity in response to *H. arabidopsidis* infection. The images in the lower panel are magnifications of the middle images. Red asterisks indicate locations where *H. arabidopsidis* haustoria formed in leaf mesophyll cells. Scale bars = 40 μm.

### Identifying host genes whose overexpression confers resistance to downy mildew

To assess whether these five genes are involved in the Arabidopsis-*H. arabidopsidis* interaction, we created Arabidopsis transformants overexpressing each gene. Two independent lines for each gene were selected. All individuals were morphologically similar to Col-0 wild-type (WT) plants (Fig. S4). At 5 d after inoculation with *H. arabidopsidis*, resistance levels of the transformants were assessed by counting the number of conidiospores formed on the plants and comparing them with Col-0 WT (Fig. 6). The most significantly resistant phenotypes were observed in *AZI3*-overexpressing lines that reproducibly had fewer than 15% of the conidiospores formed on Col-0 WT. The other resistant lines were *KTI4* overexpressors that had fewer than one-half of the conidiospores formed on Col-0 WT. Plants overexpressing *AT1G09932* appeared to have slightly increased resistance to *H. arabidopsidis*. In contrast, *PCR2*-overexpressing and *GLP9*-overexpressing lines showed no difference in resistance compared to Col-0 WT. Notably, none of the tested transformants differed from Col-0 WT in their resistance to the bacterial pathogen *Pseudomonas syringae* pv. *tomato* (*Pto*) DC3000 (Fig. 6), suggesting that at least the *AZI3*- and *KTI4*-overexpressing lines are specifically resistant to *H. arabidopsidis*. To investigate the effect of *azi3* loss on disease resistance, we searched for the available T-DNA mutants but did not find any insertions in *AZI3*; however, we did find a line with T-DNA inserted in the promoter region of *KTI4* (SALK_131716C, refer to *kti4.1*), leading to reduced *KTI4* expression (Arnaiz et al., 2018). No significant differences in disease resistance to *H. arabidopsidis* were observed for *kti4.1* compared to Col-0 WT (Fig. S6).

**Figure 6.**
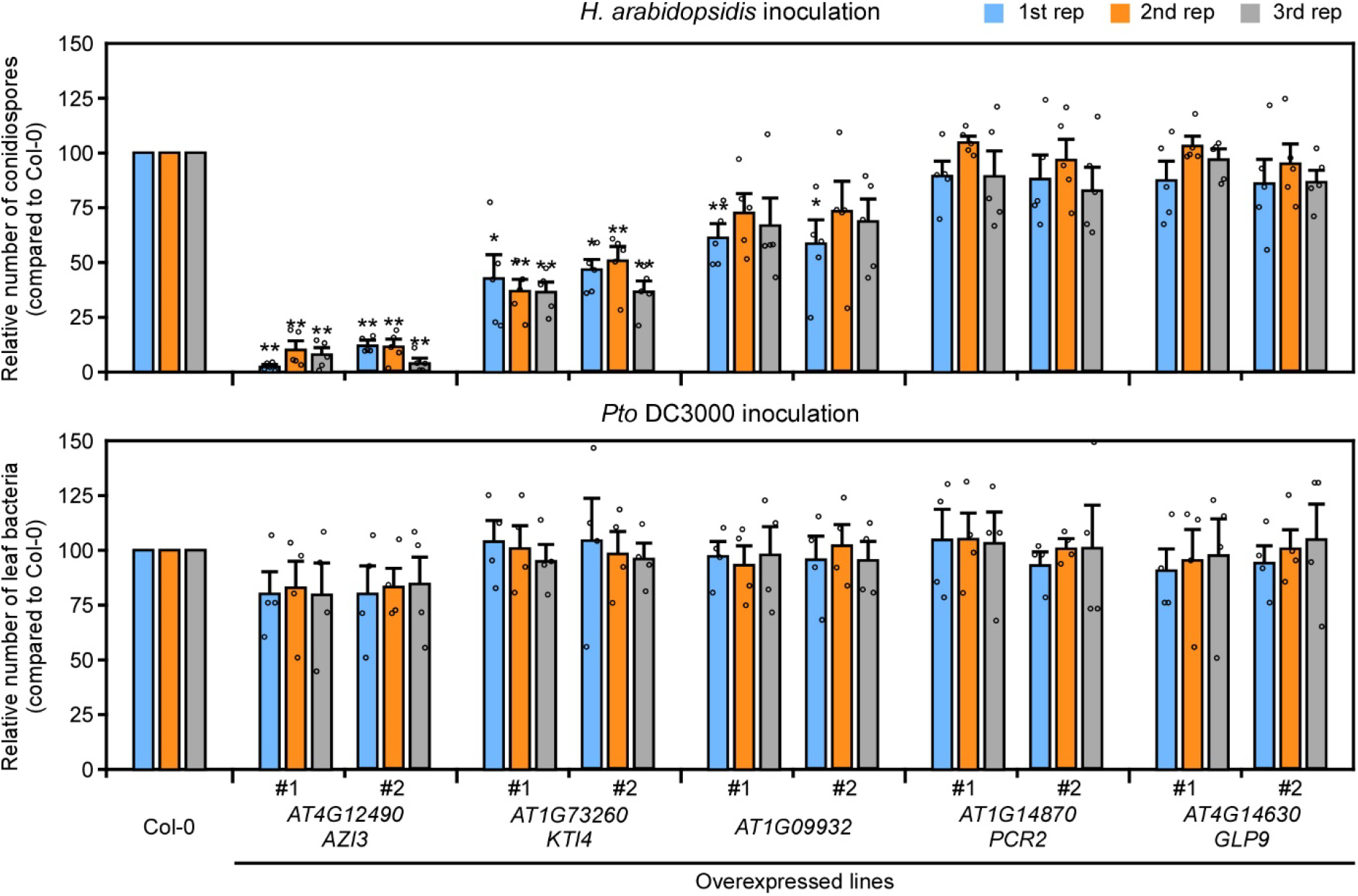
Disease resistance phenotypes of transgenic plants expressing *DMR6*-coexpressed genes. *H. arabidopsidis* (upper panel) and *P. syringae* pv. *tomato* (*Pto*) DC3000 (lower panel) growth on two independent transgenic lines expressing the indicated genes. Data are shown relative to the Arabidopsis Col-0 WT value of 100. Data are means ± SEs from five and four biological replicates for *H. arabidopsidis* and *Pto* DC3000 growth, respectively, and represent three independent results. Data were analyzed by Student’s *t*-test: *,*p* < 0.05; **, *p* < 0.01 vs Col-0 WT plants.

## Discussion

RNA profiling is a powerful method for determining the molecular basis of host-pathogen interactions, but analyses using whole tissues lead to responses from a variety of cell types, including infected- and non-infected cells. Here, we present an infected cell-specific RNA profiling strategy during the Arabidopsis/downy mildew interaction by employing a new TRAP system using the E9-Im9 pair. Our study found genes that are specifically expressed in cells haustoriated by *H. arabidopsidis*. For example, this method detected *PSK4* and *WRKY18* that are specifically expressed in haustoriated cells. Furthermore, overexpression of *AZI3* and *KTI4*, two genes found to be specifically expressed in haustoriated cells, conferred resistance to *H. arabidopsidis* but not to *Pto* DC3000.

Recently, a conceptually similar methodology using split GFPs was reported (Dinkeloo et al., 2022). Like ours, their method employed the *DMR6* promoter to drive the expression of a GFP fragment with a purification tag and another GFP fragment with a ribosome binding site, enabling the capture of polysomes from infected cells. Unfortunately, the report did not provide a list of genes detected by this method, making it impossible to compare with our dataset. One notable strategic difference is that we also used the *PR1* promoter, which is active in neighboring cells but not in haustoriated cells (Caillaud et al., 2013), to remove genes expressed in both cell types. This strategy provided an essential step as 2,953 out of 4,524 genes (65%) that *pDMR6::E9-HF* captured were also found by *pPR1::E9-HF* (Fig. 4B). Furthermore, 54 out of 1,571 (3.4%) genes were selected as induced at 5 dpi with *H. arabidopsidis* to eliminate genes expressed in haustoriated cells but not responsive to the pathogen (Fig. 4B). Finally, histochemical GUS analysis confirmed that at least 7 genes were specifically expressed in the haustoriated cells (Fig. 5). These results strongly support that our RNA profiling of the cells of interest was successful.

Among the 7 genes, we found *PSK4* and *WRKY18* that are known to be involved in the modulation of plant immunity. Overexpression of *PSK4* and application of its active 5-amino-acid bisulfated phytosulfokine (PSK) peptide inhibit pattern-triggered immunity (PTI) responses and increase the susceptibility to pathogens (Igarashi et al., 2012; Mosher et al., 2013). Similarly, *WRKY18* is redundant with *WRKY40* and negatively regulates the expression of PTI-responsive genes and resistance toward *Pto* DC3000 and the powdery mildew fungus *Golovinomyces orontii* (Xu et al., 2006; Pandey et al., 2010; Birkenbihl et al., 2017). As *PSK4* and *WRKY18* are specifically induced in haustoriated cells, these genes can be considered as *S* genes that help pathogen infection, similar to *DMR6*. A previous chromatin immunoprecipitation sequencing (ChIP-seq) analysis reported 1,403 genes as WRKY18 target genes (Birkenbihl et al., 2017). In our experiments, 9 out of the 54 genes (17%), including *DMR6* and the 53 *DMR6*-coexpressed genes, were identified (Table 1) as targets of WRKY18 (Table S4). Thus, WRKY18 may play a key role as a transcriptional hub for the *S* genes network. Since many *H. arabidopsidis* effectors are known to localize into plant cell nuclei when expressed *in planta* (Caillaud et al., 2012), targeting such hubs can be a suitable strategy for establishing infections as a biotroph.

In this study, we also identified *AZI3* as a transcriptionally induced gene in haustoriated cells whose overexpression conferred resistance to *H. arabidopsidis*. *AZI3 (AT4G12490*) is a close paralog of the lipid transfer proteins *AZI1* (*AT4G12470*) and *AZI4* (*AT4G12500*). These three genes have another paralog, *EARLY ARABIDOPSIS ALUMINUM INDUCED 1* (*EARLI1; AT4G12480*); all four genes are tandemly located on chromosome 4 in Arabidopsis (Cecchini et al., 2015), and all four genes are induced upon *H. arabidopsidis* infection (Asai et al., 2014). In particular, *EARLI1* is included among the 53 *DMR6*-coexpressed genes (Table 1), whereas *AZI1* and *AZI4* are not included but appear to be coexpressed with *DMR6* (Fig. S5). Among the four paralogs, *AZI1* and *EARLI1* are reportedly key factors in establishing systemic acquired resistance (SAR) by affecting the lipid derivative azelaic acid (AZA) mobilization from local tissues to distal sites (Jung et al., 2009; Cecchini et al., 2015). AZI1, AZI3, and EARLI1 all localize in the endoplasmic reticulum (ER)/plasmodesmata, chloroplast outer envelopes, and membrane-contact sites between these organelles (Cecchini et al., 2015). Since AZA is produced in chloroplasts (Zoeller et al., 2012), AZI1 and its paralogs are thought to form part of the complexes contacting both chloroplasts and ER membranes, potentially allowing the non-vesicular transport of AZA to distal tissues (Cecchini et al., 2015). In this scenario, Arabidopsis may induce SAR signaling to counter secondary infection by expressing *AZI1* and its paralogs in the *H. arabidopsidis-infected* cells. Consistent with this hypothesis, *AZI3* overexpressing lines exhibited enhanced resistance to *H. arabidopsidis* (Fig. 6). Interestingly, the *AZI3*-based enhanced resistance is *H. arabidopsidis* specific as *AZI3* overexpressors showed no difference in bacterial growth on local leaves after inoculation with *Pto* DC3000, a finding consistent with the results in *AZI1* overexpressing lines reported by Wang et al. (2016). The effect of *azi3* loss on disease resistance was not investigated since the corresponding T-DNA mutants were unavailable. As SAR is reduced in the *azi1* and *earli1* mutants (Jung et al., 2009; Cecchini et al., 2015), it should be instructive to determine the effect of the quadruple mutation of *AZI1* and its paralogs on plant immunity.

*KTI4*, a gene that encodes a functional Kunitz trypsin inhibitor (Li et al., 2008), is another gene identified in our study. The observation that *KTI4* overexpressors exhibit higher resistance to *H. arabidopsidis* (Fig 6) is markedly different from the findings of a previous study that reported overexpression of *KTI4* leads to higher susceptibility to the bacterial necrotroph *Pectobacterium carotovorum* (formerly *Erwinia carotovora*) (Li et al., 2008). The opposite resistance phenotypes against these pathogens might be due to a difference in lifestyle between biotrophs and necrotrophs. As SA signaling functions oppositely in biotrophs and necrotrophs (Hou and Tsuda, 2022) and *KTI4* is induced by SA (Li et al., 2008), *KTI4* may be involved in SA signaling. The expression of *DMR6* inactivates SA in *H. arabidopsidis*-haustoriated cells and may suppress plant immunity activated by *KTI4*, resulting in infection. *KTI4* overexpressing lines did not show increased resistance to *Pto* DC3000 (Fig. 6) or have any effect on plant growth (Fig. S4), suggesting that *KTI4*-mediated immunity is not constantly activated. The *kti4.1* mutant showed no difference in resistance to *H. arabidopsidis* compared to Col-0 WT (Fig. S6), possibly because of redundancy, as there are six paralogs of *KTI4* in Arabidopsis (Arnaiz et al., 2018). In fact, the closest paralog *KTI5* (*At1G17860*) seems to be coexpressed with *DMR6* (Fig. S7), although the paralog was not included in the list of 53 *DMR6*-coexpressed genes (Table 1). Further analysis is needed to determine how *KTI4* may be involved in resistance to *H. arabidopsidis*.

Our new TRAP system revealed host genes induced in the *H. arabidopsidis-infected* cells that function either in susceptibility or resistance. We hypothesize that different mechanisms induce the expression of these genes. For instance, susceptibility-related genes may be induced by *H. arabidopsidis*, perhaps by using its effectors. In contrast, Arabidopsis may actively induce resistance-related genes by recognizing pathogen-derived molecules. Further genetic analysis is needed to dissect the signaling pathways. In addition, we expect that this E9-Im9 based TRAP system could be applicable to several other stimulus-specific contexts and other plant-pathogen interactions using relevant specific promoters.

## Materials and methods

### Plant material and growth

Arabidopsis plants were grown at 22 °C with a 10-h photoperiod and a 14-h dark period in environmentally controlled growth cabinets. *N. benthamiana* plants were grown at 25 °C with a 16-h photoperiod and an 8-h dark period in environmentally controlled growth cabinets.

### Pathogen assays

Inoculation with the *H. arabidopsidis* Waco9 isolate was conducted as described by Asai et al. (2015). Briefly, Arabidopsis plants were spray-inoculated to saturation with a spore suspension of 1 x 10^4^ conidiospores/mL. Five replicates of three plants for each Arabidopsis line were used in the assays. Plants were covered with a transparent lid to maintain high humidity (90-100%) conditions in a growth cabinet at 16 °C with a 10-h photoperiod until the day of sampling. Conidiospores were harvested in 1 mL of water. After vortexing, the number of released conidiospores was determined using a hemocytometer. *P. syringae* pv. *tomato* DC3000 was grown on LB media containing 100 μg/mL rifampicin at 28 °C. Five- to six-week-old soil-grown plants were syringe-infiltrated with a bacterial suspension of 5 × 10^5^ cfu/mL in 10 mM MgCl_2_. Bacterial growth in plants was monitored at 3 d post inoculation.

### Plasmid construction

For the construction of the TRAP plasmids, the ORF of *RPL18* together with 3’ UTR and the terminator was amplified from Col-0 gDNA for Golden Gate assembly (Engler et al. 2008 PLOS One; Engler et al. 2014 ACS Synth Biol) into the pICH47751 vector with the *35S* promoter and *Im9* (with GS spacer) as an N-terminal fusion tag. The 2,486 bp *DMR6*, 2,378 bp *PR1*, and 1,450 bp *Act2* promoters were amplified from Col-0 gDNA for Golden Gate assembly (Engler et al. 2008 PLOS One; Engler et al. 2014 ACS Synth Biol) into the pICH47761 vector with *E9*, *HF* as a C-terminal fusion tag and *OCS* terminator. For the final Golden Gate assembly, *p35S::Im9-RPL18* (pICH47751) was combined with *pDMR6/pPR1/pAct2*::*E9-HF* (pICH47761), the herbicide BASTA resistance gene (*BAR*; pICH47732) and *FastRed* (pICH47742) into the Level 2 Golden Gate vector pAGM4723.

For the transient expression studies, the ORF of *RPL18* was amplified from Col-0 cDNA for Golden Gate assembly (Engler et al. 2008 PLOS One; Engler et al. 2014 ACS Synth Biol) into the binary vector pICH86988 with *Im9* or *YFP* as an N-terminal fusion tag. *E9* fused to *GFP* as a C-terminal fusion tag was also cloned into the pICH86988 vector.

For GUS reporter constructs, the promoter sequence plus 27 bp or 30 bp upstream from the start codon of *PSK4* (1,827 bp), *WRKY18* (2,030 bp), *AT1G09932* (1,062 bp), *PCR2* (2,030 bp), *KTI4* (993 bp), *AZI3* (2,030 bp) and *GLP9* (2,030 bp) was amplified from Col-0 gDNA for Golden Gate assembly (Engler et al. 2008 PLOS One; Engler et al. 2014 ACS Synth Biol) into the binary vector pICSL86955 with the *GUS* reporter gene and *OCS* terminator.

For overexpressing constructs, the ORFs of *AT1G09932, PCR2, KTI4, AZI3*, and *GLP9* were amplified from Col-0 gDNA for Golden Gate assembly (Engler et al., 2008; Engler et al., 2014) into the binary vector pICSL86977 with a C-terminal *HF* fusion tag.

### Transient gene expression and plant transformation

For transient gene expression analysis, *Agrobacterium tumefaciens* strain AGL1 was used to deliver the respective transgenes to *N. benthamiana* leaves using methods previously described (Asai et al., 2008). All bacterial suspensions carrying individual constructs were adjusted to an OD_600_ = 0.5 in the final mix for infiltration, except for the coexpression of *35S::E9-GFP* with *35S::Im9-RPL18* in which bacterial suspensions were adjusted to OD_600_ = 0.25 for *35S::E9-GFP* and OD_600_ =0.5 for *35S::Im9-RPL18* due to low expression levels of Im9-RPL18. We hypothesize that the turnover of RPL18 occurs more rapidly than for E9-GFP.

For plant transformation, Arabidopsis Col-0 plants were transformed using the dipping method (Clough and Bent, 1998). Briefly, flowering Arabidopsis plants were dipped into a solution containing *A. tumefaciens* carrying a plasmid of interest, and the seeds were harvested to select the T1 transformants on selective MS media. T1 plants were checked for expression of the construct-of-interest by immunoblot analysis. T2 seeds were sown on selective MS media, and the proportion of resistant versus susceptible plants was measured to identify lines with single T-DNA insertions. Transformed plants were transferred to soil, and the seeds were collected. Two independent T3 homozygous lines were analyzed.

### Confocal microscopy

For *in planta* subcellular localization analysis in *N. benthamiana*, cut leaf patches were mounted in water and analyzed using a Leica TCS SP8 X confocal microscope (Leica Microsystems) with the following excitation wavelengths. GFP, 488 nm; YFP, 513 nm.

### Protein extraction and immunoblotting

Leaves were ground to a fine powder in liquid nitrogen and thawed in extraction buffer (50 mM Tris-HCl, pH 7.5, 150 mM NaCl, 10% (v/v) glycerol, 10 mM DTT, 10 mM EDTA, 1 mM NaF, 1 mM Na_2_MoO_4_.2H_2_O, 1% (v/v) IGEPAL CA-630 from Sigma-Aldrich and 1% (v/v) protease inhibitor cocktail from Sigma-Aldrich). Samples were cleared by centrifugation at 16,000 *g* for 15 min at 4 °C, and the supernatant liquid was collected and subjected to SDS-PAGE. Proteins were then electroblotted onto a PVDF membrane using a semidry blotter (Trans-Blot Turbo Transfer System; Bio-Rad). Membranes were blocked overnight at 4 °C in TBS-T (50 mM Tris-HCl, pH 7.5, 150 mM NaCl, and 0.05% (v/v) Tween 20) with 5% (w/v) skim milk. Membranes were then incubated with horseradish peroxidase-conjugated anti-FLAG antibody (1:20,000; A8592; Sigma-Aldrich) diluted with TBS-T with 5% (w/v) skim milk at room temperature for 1 h. After washing with TBS-T, bound antibodies were visualized using SuperSignal West Femto Maximum Sensitivity Substrate (Thermo Fisher Scientific). Bands were imaged using an image analyzer (ImageQuant LAS 4000 imager; GE Healthcare).

### Translating ribosome affinity purification (TRAP)

TRAP was performed according to the method of Mustroph et al. (2009a) with the following modifications: 8 mL of polysome extraction buffer (PEB) was added to 81 samples of 3-week-old plant-derived tissues that were ground in liquid nitrogen. The resulting extract was clarified twice by centrifugation at 16,000 *g* for 15 min at 4 °C, with a Miracloth filtration step between centrifugations. From a portion of the clarified extract, RNA was extracted and referred to as RNAs_Total. The remainder of the extract was mixed with 150 μL washed α-FLAG agarose beads (A2220, Sigma) and adjusted to 5 mL with PEB. The extract was incubated with the beads for 2 h with gentle rocking at 4 °C. The beads were washed as follows: one wash with 6 mL PEB and four washes with 6 mL wash buffer. The washed beads were resuspended in 300 μL wash buffer containing 300 ng/μL of 3xFLAG peptide (F4799, Sigma) and 20 U/mL RNAsin (Promega) and incubated for 30 min with gentle rocking at 4 °C. RNA was extracted from the supernatant liquid collected after centrifugation and is referred to as RNAs_IP.

### RNA extraction, cDNA synthesis, and RT-qPCR

Total RNAs were extracted using RNeasy Plant Mini Kit (Qiagen) according to the manufacturer’s procedure. Total RNAs (1 μg) were used for generating cDNAs in a 20 μL reaction according to the Invitrogen Superscript III Reverse Transcriptase protocol. The obtained cDNAs were diluted five times, and 1 μL was used for a 10 μL qPCR reaction. qPCR was performed in a 10 μL final volume using 5 μL SYBR Green Mix (Toyobo), 1 μL diluted cDNAs, and primers. qPCR was run on Mx3000P qPCR System (Agilent) using the following program: (1) 95 °C, 3 min; (2) [95 °C, 30 sec, then 60 °C, 30 sec, then 72 °C, 30 sec] x 45, (3) 95 °C, 1 min followed by a temperature gradient from 55 °C to 95 °C. The relative expression values were determined using the comparative cycle threshold method (2^−ΔΔCt^). *EF-1α* was used as the reference gene. Primers used for qPCR are listed in Supplementary Table S4.

### RNA sequencing

The library prepared for RNA sequencing was constructed as described previously (Rallapalli et al., 2014). Purified double-stranded cDNAs were subjected to Covaris shearing (parameters: intensity, 5; duty cycle, 20%; cycles/burst, 200; duration, 60 sec). The libraries were sequenced on an Illumina NextSeq 500 DNA sequencer. Sequence data have been deposited in NCBI’s Gene Expression Omnibus (GEO) and are accessible through GEO Series accession number GSE220449. The Illumina sequence library was quality-filtered using FASTX Toolkit version 0.0.13.2 (Hannonlab) with parameters −q20 and −p50. Reads containing “N” were discarded. Quality-filtered libraries were aligned on the Arabidopsis Col-0 genome with the Araport11 annotation using the default settings of CLC Genomic Workbench 20. Transcription levels for each transcript were calculated as TPM (transcripts per million). Differential expression was analyzed using the R statistical language version 4.1.1 with edgeR version 3.34.0 (Robinson et al., 2010), part of the Bioconductor package (Gentleman et al., 2004). GO analysis of the 54 confident candidate *DMR6*-coexpressed genes shown in Table 1 used PANTHER (Mi et al., 2019) at The Arabidopsis Information Resource (TAIR) website (https://www.arabidopsis.org/tools/go_term_enrichment.jsp).

### GUS staining

GUS activity was assayed histochemically with 5-bromo-4-chloro-3-indolyl-β-D-glucuronic acid (1 mg/mL or 0.2 mg/mL) in a buffer containing 100 mM sodium phosphate pH 7.0, 0.5 mM potassium ferrocyanide, 0.5 mM potassium ferricyanide, 10 mM EDTA, 0.1% (v/v) Triton. Arabidopsis leaves were vacuum infiltrated with staining solution and then incubated overnight at 37 °C in the dark. Samples were destained in absolute ethanol followed by incubation in a chloral hydrate solution. Stained leaves were observed using an Olympus BX51 microscope.

## Supplemental data

The following materials are available in the online version of this article.

**Supplemental Figure S1. Schematic diagram of the traditional translating ribosome affinity purification (TRAP) system.** (**A**) A schematic representation of the chimeric construct; an epitope-tagged ribosomal protein L18 (RPL18) fused to a constitutive promoter or a promoter that is active in a specific cell type (cell-type promoter). (**B**) Schematic diagram of ribosomal complexes in cells where the promoters fused to an epitope-tagged RPL18 are unactivated or activated.

**Supplemental Figure S2. Expression levels of *DMR6* and *PR1* in samples derived from whole tissues during *H. arabidopsidis* infection.** Expression levels of *DMR6* (**A**) and *PR1* (**B**) at 1, 3, and 5 d post inoculation (dpi) with *H. arabidopsidis* Waco9 isolate or water as a control (Mock) are represented as TPM (tags per million) of total reads aligned to the Arabidopsis genome. The data are derived from RNA seq data from Asai et al. 2014.

**Supplemental Figure S3. Enriched gene ontology (GO) terms of *DMR6*-coexpressed genes.** GO enrichment analysis of *DMR6* and 53 *DMR6*-coexpressed genes. Fold enrichment (*p* < 0.05) was determined by query gene number divided by the expected gene number for each GO term.

**Supplemental Figure S4. Morphology of transgenic plants overexpressing *DMR6*-coexpressed genes.** (**A**) Confirmation of protein accumulation in Arabidopsis Col-0 transgenic lines overexpressing *AT1G09932, PCR2, KTI4, AZI3*, and *GLP9*. Total proteins were prepared from 6-week-old plants. Immunoblotted proteins were treated with anti-FLAG (upper panel) antibodies. Protein loads were monitored by Coomassie Brilliant Blue (CBB) staining of the bands corresponding to ribulose-1,5-bisphosphate carboxylase (Rubisco) large subunit (lower panel). (**B**) Morphology of Arabidopsis Col-0 WT and transgenic lines photographed at 6-weeks.

**Supplemental Figure S5. Expression levels of *AZI1, EARLI1, AZI3*, and *AZI4* in the TRAP samples.** Expression of *AZI1* (*AT4G12470*), *EARLI1* (*AT4G12480*), *AZI3* (*AT4G12490*) and *AZI4* (*AT4G12500*) in the Arabidopsis Col-0 transgenic lines containing *pDMR6::E9-HF* (pDMR6), *pPR1::E9-HF* (pPR1) or *pAct2::E9-HF* (pAct2) and *p35S::Im9-RLP18*. The expression level of *AZI1, EARLI1, AZI3*, and *AZI4* in the RNAs_Total and RNAs_IP samples are represented as TPM (transcripts per million) of total reads aligned to the Arabidopsis genome. Data are means ± SDs from three biological replicates.

**Supplemental Figure S6. Disease resistance phenotypes of *kti4.1* to *H. arabidopsidis* inoculation.** *H. arabidopsidis* growth (conidiospore number) on the *kti4.1* mutant. Data are shown relative to the Arabidopsis Col-0 WT value of 100. Data are means ± SEs from five biological replicates and represent three independent results.

**Supplemental Figure S7. Expression levels of *KTI4* and its paralogs in the TRAP samples.** Expression of *KTI4* and its paralogs in the Arabidopsis Col-0 transgenic lines containing *pDMR6::E9-HF* (pDMR6), *pPR1::E9-HF* (pPR1) or *pAct2::E9-HF* (pAct2) and *p35S::Im9-RLP18*. The expression level of *KTI4* and its paralogs in RNAs_Total and RNAs_IP samples are represented as TPM (transcripts per million) of total reads aligned to the Arabidopsis genome. Data are means ± SDs from three biological replicates. ND, not detectable.

**Supplemental Table S1. The expression patterns of Arabidopsis genes in the TRAP samples after inoculation with *H. arabidopsidis* Waco9.**

**Supplemental Table S2. Genes with significantly different expression levels.**

**Supplemental Table S3. Gene accession numbers for differentially expressed candidate genes.**

**Supplemental Table S4. Primers used in this study.**

## Acknowledgments

We thank Dr. Sylvestre Marillonnet for Golden Gate vectors and Prof Colin Kleanthous (U Oxford) for helpful discussion about the E9/Im9 system and for providing E9 and Im9 clones. We are grateful to Ryo Yoshida for providing the illustrations. We also thank Takuya Okubo, Soshi Tsuchiya, Asuka Yoshida, Kota Hidaka, Manami Yamazaki, Ippei Takahashi, Eri Kurai, Emika Okubo, and Kaoru Yoshida for their support.

## Author Contributions

S.A., V.C., J.D.G.J. and K.S. conceptualized and designed the research. S.A. and V.C. conducted experiments and data analysis. S.A. and K.S. wrote the manuscript.

## References

Arnaiz A, Talavera-Mateo L, Gonzalez-Melendi P, Martinez M, Diaz I, Santamaria ME (2018) Arabidopsis kunitz trypsin inhibitors in defense against spider mites. Front Plant Sci 9: 986

Asai S, Ohta K, Yoshioka H (2008) MAPK signaling regulates nitric oxide and NADPH oxidase-dependent oxidative bursts in *Nicotiana benthamiana*. Plant Cell 20: 1390–1406

Asai S, Rallapalli G, Piquerez SJ, Caillaud MC, Furzer OJ, Ishaque N, Wirthmueller L, Fabro G, Shirasu K, Jones JD (2014) Expression profiling during Arabidopsis/downy mildew interaction reveals a highly-expressed effector that attenuates responses to salicylic acid. PLoS Pathog 10: e1004443

Asai S, Shirasu K, Jones JD (2015) *Hyaloperonospora arabidopsidis* (downy mildew) infection assay in *Arabidopsis*. Bio-protocol 5: e1627

Birkenbihl RP, Kracher B, Roccaro M, Somssich IE (2017) Induced genome-wide binding of three Arabidopsis WRKY transcription factors during early MAMP-triggered immunity. Plant Cell 29: 20–38

Caillaud MC, Asai S, Rallapalli G, Piquerez SJM, Fabro G, Jones JDG (2013) A downy mildew effector attenuates salicylic acid-triggered immunity in Arabidopsis by interacting with the host mediator complex. PLoS Biol 11: e1001732

Cecchini NM, Steffes K, Schlappi MR, Gifford AN, Greenberg JT (2015) *Arabidopsis* AZI1 family proteins mediate signal mobilization for systemic defence priming. Nat Commun 6: 7658

Clough SJ, Bent AF (1998) Floral dip: a simplified method for *Agrobacterium*-mediated transformation of *Arabidopsis thaliana*. Plant J 16: 735–743

Coates ME, Beynon JL (2010) *Hyaloperonospora Arabidopsidis* as a pathogen model. Annu Rev Phytopathol 48: 329–345

Dinkeloo K, Pelly Z, McDowell JM, Pilot G (2022) A split green fluorescent protein system to enhance spatial and temporal sensitivity of translating ribosome affinity purification. Plant J 111: 304–315

Engler C, Kandzia R, Marillonnet S (2008) A one pot, one step, precision cloning method with high throughput capability. PLoS One 3: e3647

Engler C, Youles M, Gruetzner R, Ehnert TM, Werner S, Jones JD, Patron NJ, Marillonnet S (2014) A golden gate modular cloning toolbox for plants. ACS Synth Biol 3: 839–843

Gentleman RC, Carey VJ, Bates DM, Bolstad B, Dettling M, Dudoit S, Ellis B, Gautier L, Ge Y, Gentry J, Hornik K, Hothorn T, Huber W, Iacus S, Irizarry R, Leisch F, Li C, Maechler M, Rossini AJ, Sawitzki G, Smith C, Smyth G, Tierney L, Yang JY, Zhang J (2004) Bioconductor: open software development for computational biology and bioinformatics. Genome Biol 5: R80

Heiman M, Kulicke R, Fenster RJ, Greengard P, Heintz N (2014) Cell type-specific mRNA purification by translating ribosome affinity purification (TRAP). Nat Protoc 9: 1282–1291

Hou S, Tsuda K (2022) Salicylic acid and jasmonic acid crosstalk in plant immunity. Essays Biochem 66: 647–656

Igarashi D, Tsuda K, Katagiri F (2012) The peptide growth factor, phytosulfokine, attenuates pattern-triggered immunity. Plant J 71: 194–204

Jung HW, Tschaplinski TJ, Wang L, Glazebrook J, Greenberg JT (2009) Priming in systemic plant immunity. Science 324: 89–91

Li J, Brader G, Palva ET (2008) Kunitz trypsin inhibitor: an antagonist of cell death triggered by phytopathogens and fumonisin b1 in *Arabidopsis*. Mol Plant 1: 482–495

Li W, Dennis CA, Moore GR, James R, Kleanthous C (1997) Protein-protein interaction specificity of Im9 for the endonuclease toxin colicin E9 defined by homologue-scanning mutagenesis. J Biol Chem 272: 22253–22258

Mi H, Muruganujan A, Ebert D, Huang X, Thomas PD (2019) PANTHER version 14: more genomes, a new PANTHER GO-slim and improvements in enrichment analysis tools. Nucleic Acids Res 47: D419–D426

Mosher S, Seybold H, Rodriguez P, Stahl M, Davies KA, Dayaratne S, Morillo SA, Wierzba M, Favery B, Keller H, Tax FE, Kemmerling B (2013) The tyrosine-sulfated peptide receptors PSKR1 and PSY1R modify the immunity of Arabidopsis to biotrophic and necrotrophic pathogens in an antagonistic manner. Plant J 73: 469–482

Mustroph A, Juntawong P, Bailey-Serres J (2009a) Isolation of plant polysomal mRNA by differential centrifugation and ribosome immunopurification methods. Methods Mol Biol 553: 109–126

Mustroph A, Zanetti ME, Jang CJ, Holtan HE, Repetti PP, Galbraith DW, Girke T, Bailey-Serres J (2009b) Profiling translatomes of discrete cell populations resolves altered cellular priorities during hypoxia in *Arabidopsis*. Proc Natl Acad Sci U S A 106: 18843–18848

Pandey SP, Roccaro M, Schon M, Logemann E, Somssich IE (2010) Transcriptional reprogramming regulated by WRKY18 and WRKY40 facilitates powdery mildew infection of Arabidopsis. Plant J 64: 912–923

Rallapalli G, Kemen EM, Robert-Seilaniantz A, Segonzac C, Etherington GJ, Sohn KH, MacLean D, Jones JD (2014) EXPRSS: an Illumina based high-throughput expression-profiling method to reveal transcriptional dynamics. BMC Genomics 15: 341

Robinson MD, McCarthy DJ, Smyth GK (2010) edgeR: a Bioconductor package for differential expression analysis of digital gene expression data. Bioinformatics 26: 139–140

van Damme M, Huibers RP, Elberse J, Van den Ackerveken G (2008) Arabidopsis *DMR6* encodes a putative 2OG-Fe(II) oxygenase that is defense-associated but required for susceptibility to downy mildew. Plant J 54: 785–793

van Schie CC, Takken FL (2014) Susceptibility genes 101: how to be a good host. Annu Rev Phytopathol 52: 551–581

Wang XY, Li DZ, Li Q, Ma YQ, Yao JW, Huang X, Xu ZQ (2016) Metabolomic analysis reveals the relationship between *AZI1* and sugar signaling in systemic acquired resistance of Arabidopsis. Plant Physiol Biochem 107: 273–287

Wirthmueller L, Asai S, Rallapalli G, Sklenar J, Fabro G, Kim DS, Lintermann R, Jaspers P, Wrzaczek M, Kangasjaervi J, MacLean D, Menke FLH, Banfield MJ, Jones JDG (2018) Arabidopsis downy mildew effector HaRxL106 suppresses plant immunity by binding to RADICAL-INDUCED CELL DEATH1. New Phytologist 220: 232–248

Xu X, Chen C, Fan B, Chen Z (2006) Physical and functional interactions between pathogen-induced Arabidopsis WRKY18, WRKY40, and WRKY60 transcription factors. Plant Cell 18: 1310–1326

Zhang Y, Zhao L, Zhao J, Li Y, Wang J, Guo R, Gan S, Liu CJ, Zhang K (2017) *S5H/DMR6* Encodes a Salicylic Acid 5-Hydroxylase That Fine-Tunes Salicylic Acid Homeostasis. Plant Physiol 175: 1082–1093

Zoeller M, Stingl N, Krischke M, Fekete A, Waller F, Berger S, Mueller MJ (2012) Lipid profiling of the Arabidopsis hypersensitive response reveals specific lipid peroxidation and fragmentation processes: biogenesis of pimelic and azelaic acid. Plant Physiol 160: 365–378

