## Supplemental Figures for "Inducible TRAP RNA profiling reveals host genes expressed in Arabidopsis cells haustoriated by downy mildew"

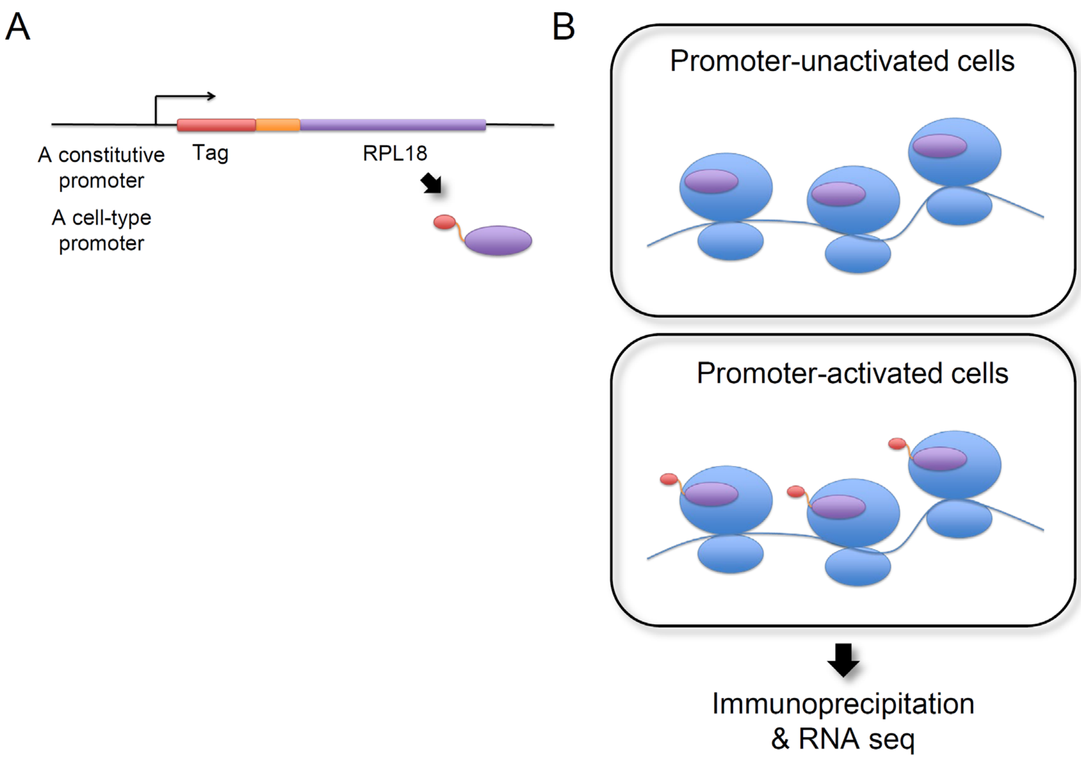


**Supplemental Figure S1. Schematic diagram of the traditional translating ribosome affinity purification (TRAP) system.** (**A**) A schematic representation of the chimeric construct; an epitope-tagged ribosomal protein L18 (RPL18) fused to a constitutive promoter or a promoter that is active in a specific cell type (cell-type promoter). (**B**) Schematic diagram of ribosomal complexes in cells where the promoters fused to an epitope-tagged RPL18 are unactivated or activated.

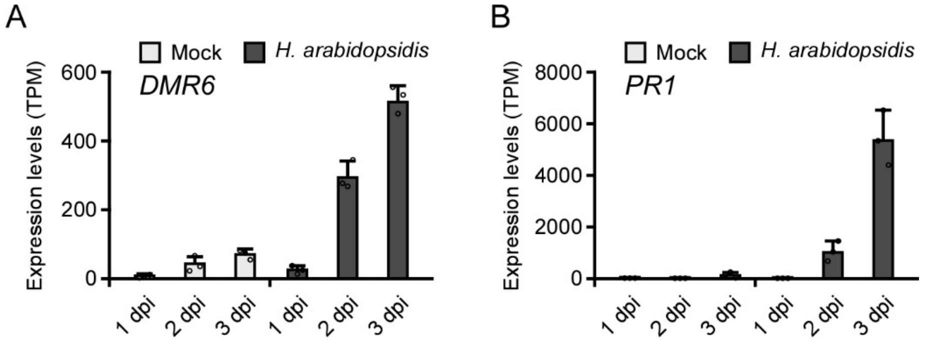


**Supplemental Figure S2. Expression levels of *DMR6* and *PR1* in samples derived from whole tissues during *H. arabidopsidis* infection.** Expression levels of *DMR6* (**A**) and *PR1* (**B**) at 1, 3, and 5 d post inoculation (dpi) with *H. arabidopsidis* Waco9 isolate or water as a control (Mock) are represented as TPM (tags per million) of total reads aligned to the Arabidopsis genome. The data are derived from RNA seq data from Asai et al. 2014.

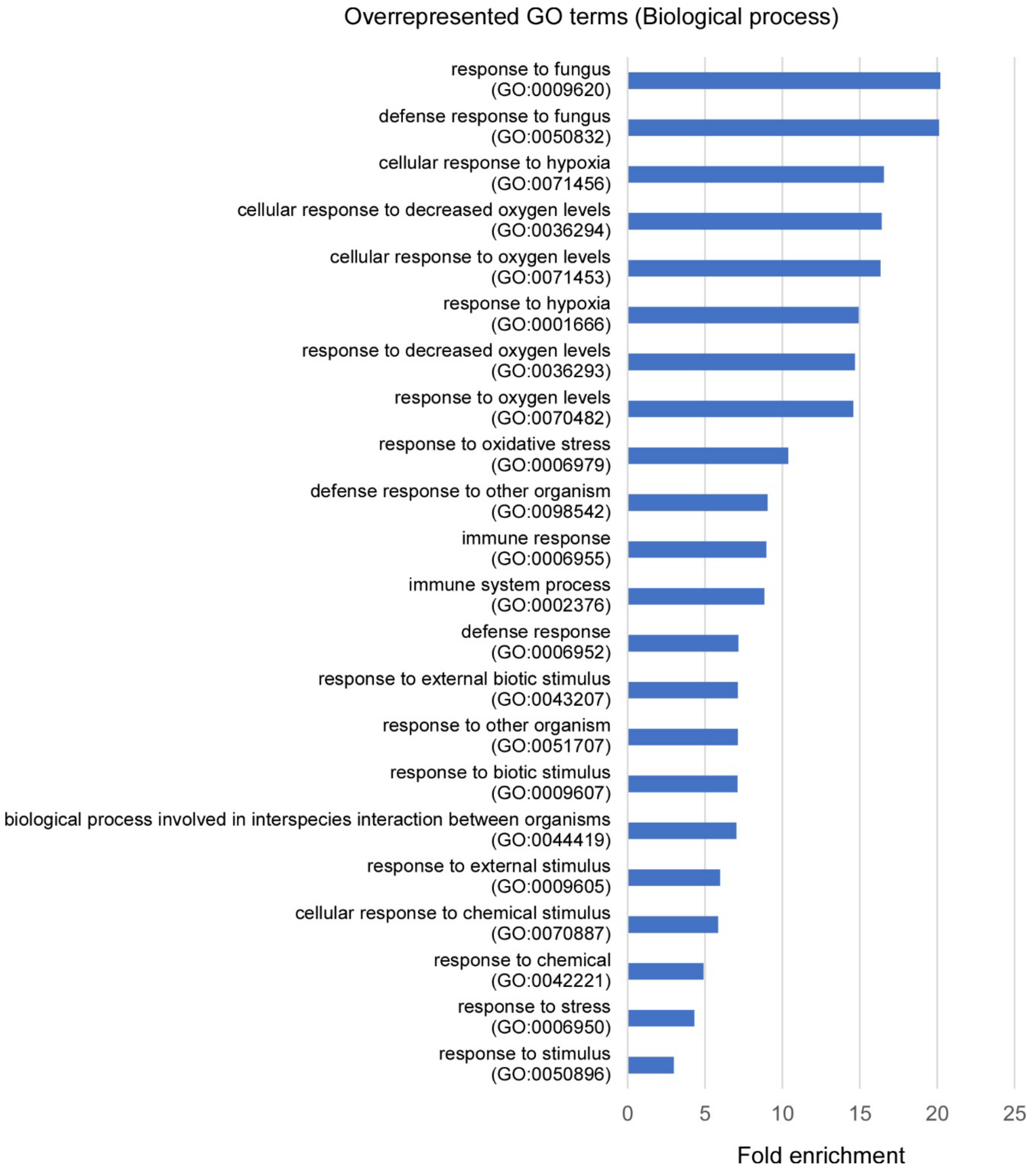


**Supplemental Figure S3. Enriched gene ontology (GO) terms of *DMR6*-coexpressed genes.** GO enrichment analysis of *DMR6* and 53 *DMR6*-coexpressed genes. Fold enrichment ($p < 0.05$) was determined by query gene number divided by the expected gene number for each GO term.

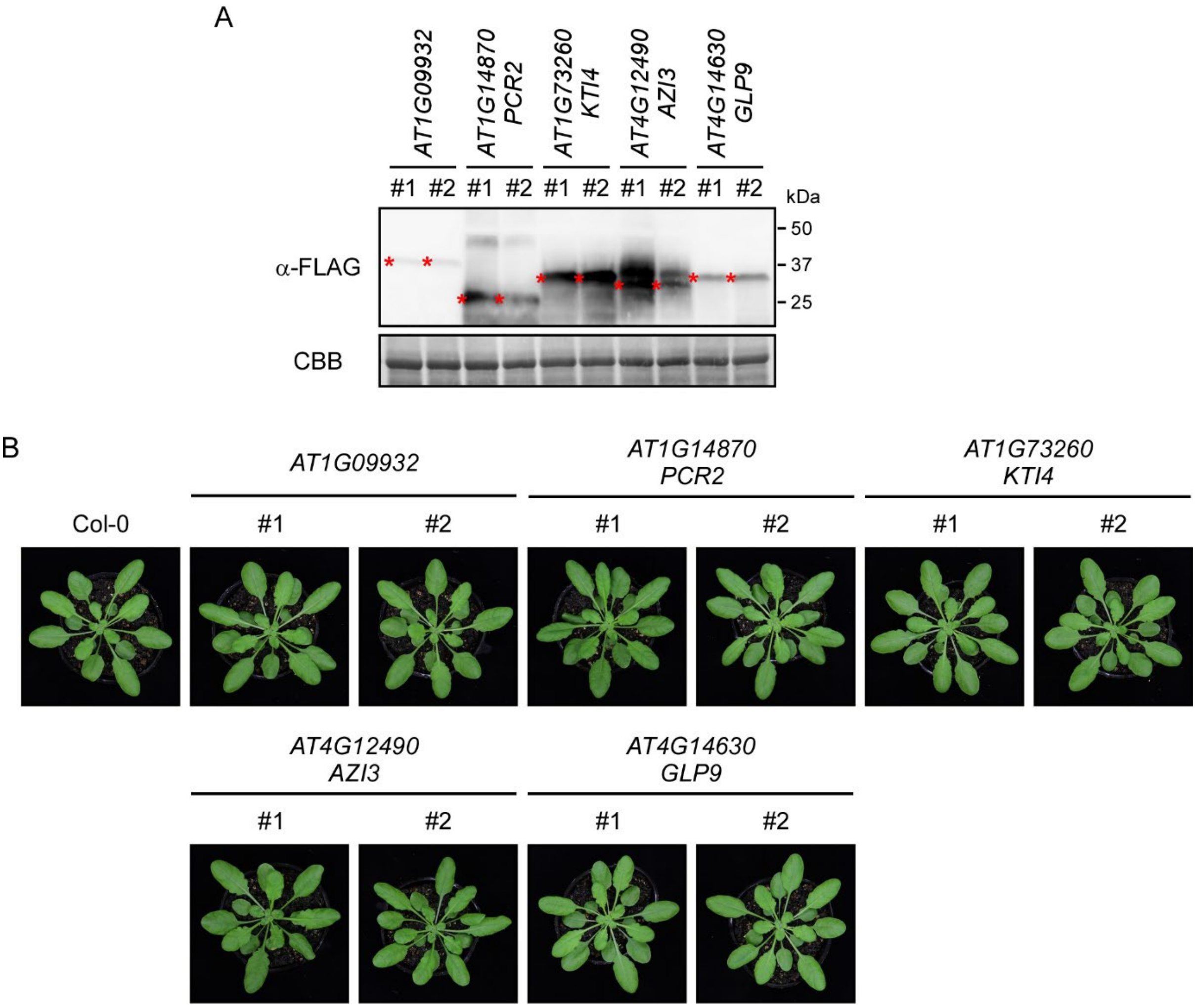


**Supplemental Figure S4. Morphology of transgenic plants overexpressing *DMR6*-coexpressed genes.** (**A**) Confirmation of protein accumulation in Arabidopsis Col-0 transgenic lines overexpressing *AT1G09932*, *PCR2*, *KTI4*, *AZI3*, and *GLP9*. Total proteins were prepared from 6-week-old plants. Immunoblotted proteins were treated with anti-FLAG (upper panel) antibodies. Protein loads were monitored by Coomassie Brilliant Blue (CBB) staining of the bands corresponding to ribulose-1,5-bisphosphate carboxylase (Rubisco) large subunit (lower panel). (**B**) Morphology of Arabidopsis Col-0 WT and transgenic lines photographed at 6-weeks.

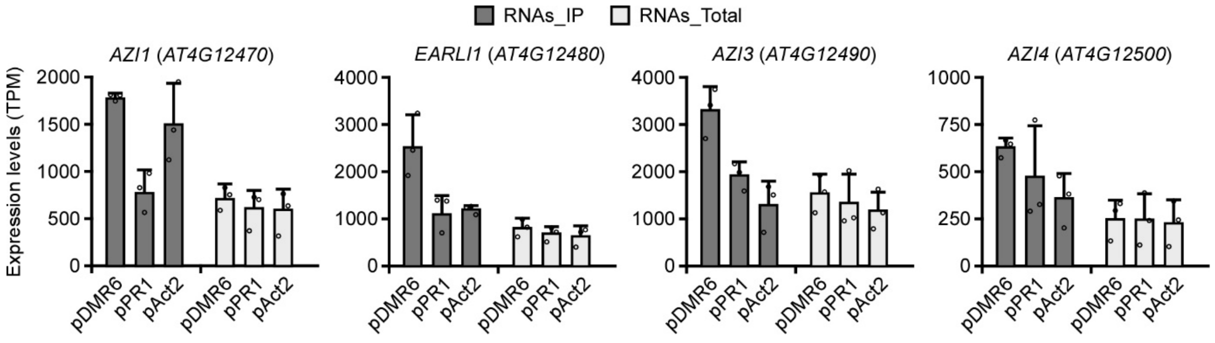


**Supplemental Figure S5. Expression levels of *AZI1*, *EARLI1*, *AZI3,* and *AZI4* in the TRAP samples.** Expression of *AZI1* (*AT4G12470*), *EARLI1* (*AT4G12480*), *AZI3* (*AT4G12490*) and *AZI4* (*AT4G12500*) in the Arabidopsis Col-0 transgenic lines containing *pDMR6*::*E9-HF* (pDMR6), *pPR1*::*E9-HF* (pPR1) or *pAct2*::*E9-HF* (pAct2) and *p35S*::*Im9-RLP18*. The expression level of *AZI1*, *EARLI1*, *AZI3,* and *AZI4* in the RNAs_Total and RNAs_IP samples are represented as TPM (transcripts per million) of total reads aligned to the Arabidopsis genome. Data are means ± SDs from three biological replicates.

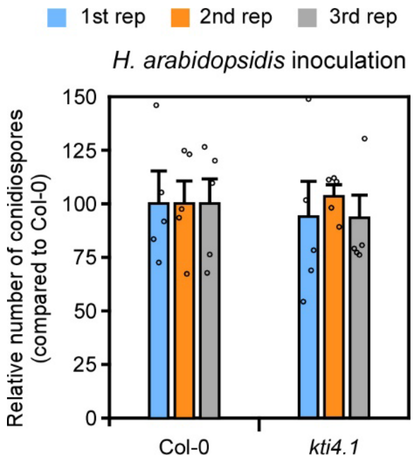


**Supplemental Figure S6. Disease resistance phenotypes of *kti4.1* to *H. arabidopsidis* inoculation.** *H. arabidopsidis* growth (conidiospore number) on the *kti4.1* mutant. Data are shown relative to the Arabidopsis Col-0 WT value of 100. Data are means ± SEs from five biological replicates and represent three independent results.

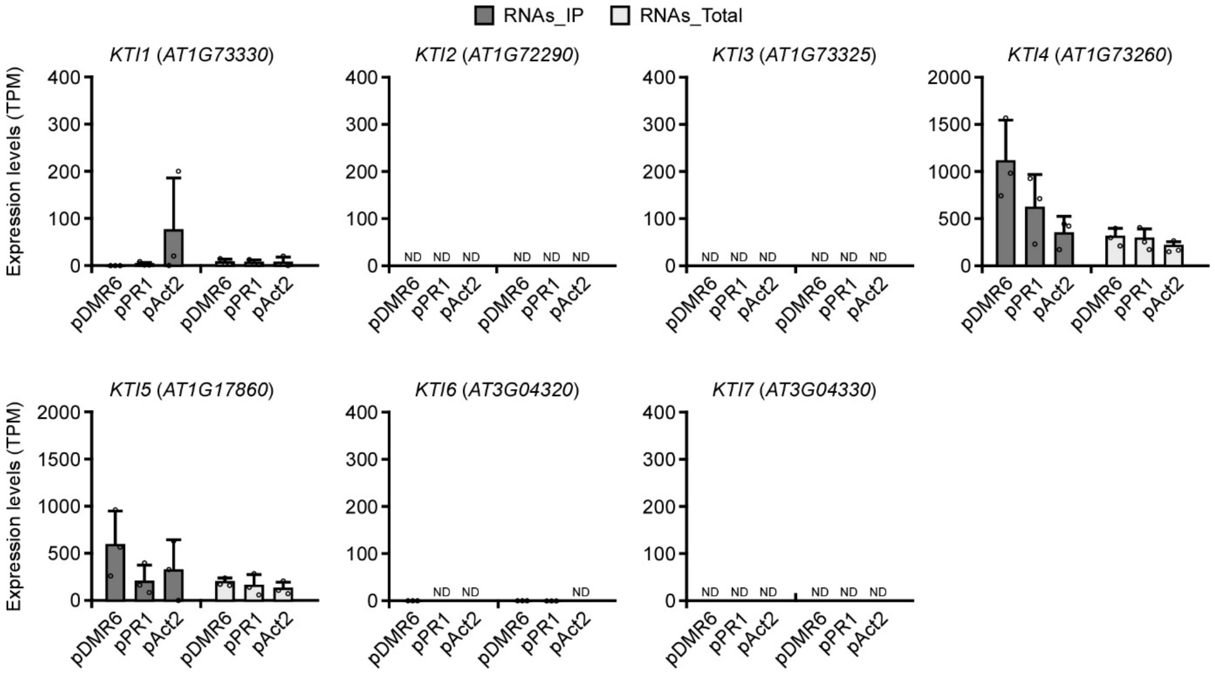


**Supplemental Figure S7. Expression levels of *KTI4* and its paralogs in the TRAP samples.** Expression of *KTI4* and its paralogs in the Arabidopsis Col-0 transgenic lines containing *pDMR6*::*E9-HF* (pDMR6), *pPR1*::*E9-HF* (pPR1) or *pAct2*::*E9-HF* (pAct2) and *p35S*::*Im9-RLP18*. The expression level of *KTI4* and its paralogs in RNAs_Total and RNAs_IP samples are represented as TPM (transcripts per million) of total reads aligned to the Arabidopsis genome. Data are means ± SDs from three biological replicates. ND, not detectable.
